# Calcium ion chelation preserves platelet function during cold storage

**DOI:** 10.1101/2020.06.14.150920

**Authors:** Binggang Xiang, Guoying Zhang, Yan Zhang, Congqing Wu, Smita Joshi, Andrew J. Morris, Jerry Ware, Susan S. Smyth, Sidney W. Whiteheart, Zhenyu Li

## Abstract

**Objective:** Platelet transfusion is a life-saving therapy to prevent or treat bleeding in patients with thrombocytopenia or platelet dysfunction. However, for more than six decades, safe and effective strategies for platelet storage have been an impediment to widespread use of platelet transfusion. Refrigerated platelets are cleared rapidly from circulation, precluding cold storage of platelets for transfusion. Consequently, platelets are stored at room temperature (RT) with an upper limit of 5 days due to risks of bacterial contamination and loss of platelet function. This practice severely limits platelet availability for transfusion. This study is to identify the mechanism of platelet clearance after cold storage and develop a method for platelet cold storage.

**Approach and Results:** We found that rapid clearance of cold-stored platelets was largely due to integrin activation and apoptosis. Deficiency of integrin β3 or caspase-3 prolonged cold-stored platelets in circulation. Pre-treatment of platelets with EGTA, a cell impermeable calcium ion chelator, reversely inhibited cold storage-induced platelet activation and consequently prolonged circulation of cold-stored platelets. Moreover, transfusion of EGTA-treated, cold-stored platelets, but not RT-stored platelets, into the mice deficient in glycoprotein Ibα significantly shortened tail-bleeding times and diminished blood loss.

**Conclusion:** Integrin activation and apoptosis is the underlying mechanism of rapid clearance of platelets after cold storage. Addition of a cell impermeable calcium ion chelator to platelet products is potentially a simple and effective method to enable cold storage of platelets for transfusion.

## Introduction

Platelets play an essential role in hemostasis. The primary physiological function of platelets is to form hemostatic thrombi that prevent blood loss and maintain vascular integrity. Platelet transfusion has been widely used to treat thrombocytopenia and platelet dysfunction and also in patients receiving myelosuppressive chemotherapy. Approximately 2.2 million platelet products are transfused each year in the United States ^1^. Unfortunately, current approaches for platelet storage limit the availability of platelets for transfusion. Like red blood cells, platelets for transfusion were originally refrigerated ^2^. In the late 1960’s, several studies reported that chilled platelets were cleared rapidly from circulation after transfusion ^2-4^, which lead to abandonment of cold storage. Since then, platelet products for transfusion are routinely stored at 20°C to 24°C with gentle agitation. Room temperature storage increases the risk of bacterial contamination. 1 in 1000 to 2000 platelet products is contaminated by bacteria, which could cause sepsis after transfusion ^5^. RT storage also leads to loss of platelet hemostatic function due to increased platelet metabolism ^3^. The current shelf life of platelet products is limited to 5 days and the bacterial testing is required ^6, 7^. As a result, about 25% of platelets collected for transfusion are not used.

Cold storage of platelets can minimize growth of contaminating bacteria and reduce platelet metabolism, thus prolonging the storage time and reducing waste. Indeed, cold storage has been shown to decrease lactate accumulation ^8^ and better preserve its aggregation response *in vitro* compared with RT storage ^3, 8-10^. The major limitation of cold storage of platelets is that chilled platelets are cleared rapidly from circulation after transfusion ^2-4, 11^. Cold storage induces platelets to change in morphology from discoid to spherical shape ^12, 13^. Cold storage also induces an increase in intracellular Ca^2+^ concentration ^14^ and cytoskeletal rearrangements ^15^. Spontaneous aggregation occurs in the cold stored platelets when they are warmed and stirred at 37°C ^16^. Cold storage also primes platelets and enhances platelet aggregation in response to subsequent agonist stimulation ^3^. However, whether those changes contribute to platelet clearance after cold storage remains unknown. Exposure of the N-acetyl-glucosamine terminals on GPIbα and clustering of GPIbα is thought to be the major mechanisms causing cold-induced platelet clearance ^17-20^. Clustered GPIbα is recognized by integrin αMβ2 on macrophages leading to platelet clearance by macrophages, mainly in liver. While short-term storage at low temperature leads to exposure of the N-acetyl-glucosamine terminals on GPIbα, after long-term storage in the cold, platelets lose sialic acid by the sialidases, Neu1 and Neu3 ^21^. Sialic acid-deprived platelets are cleared by hepatocytes mediated by the Ashwell-Morell receptor ^20, 22, 23^. A clinical trial attempting to restore survival of chilled platelets by increasing GPIbα glycosylation failed, suggesting that increased clearance after cold storage is not solely due to deglycosyalation of GPIbα. In the present study, we showed that cold storage induced integrin activation, which is responsible for cold-induced rapid clearance. Cold-stored platelets from integrin β3 deficient mice remained in circulation, similar as observed with fresh platelets, for at least the first 8 hours after transfusion. A cell impermeable calcium ion chelator EGTA reversibly inhibited cold temperature-induced integrin activation and aggregation and prevented rapid clearance after cold storage. Our data reveal a novel mechanism of cold storage-induced platelet clearance and also indicate that inclusion of cell impermeable calcium chelators can prevent platelet from rapid clearance after cold storage.

## Results

### RT-stored platelets were cleared rapidly after transfusion

Previous studies used labeled platelets, either with radioactive material or fluorescent dye, to differentiate transfused platelets from endogenous (ie recipient) platelets ^2, 17^. It is unknown whether the labeling process might affect platelet structure and life span after transfusion. To minimize the possible effects of manipulation of platelets before transfusion, we developed a novel model to monitor platelet life span in circulation after transfusion. Platelets from C57BL/6-Tg(CAGEGFP)1Osb/J mice are GFP positive and are distinguishable from wild-type recipient platelets by flow cytometry assay (**Fig. 1A**). Injection of 2.5 × 10^8^ GFP-positive fresh platelets into a ∼20 g mouse resulted in ∼15% GFP-positive platelet population. Consistent with previous findings, the life span of mouse platelets determined using this method is 4∼5 days (**Fig. 1A** and **1B**). We then evaluated life span of RT-stored platelets in circulation by monitoring the ratio of transfused (GFP-positive) platelets versus recipient (GFP-negative) platelets. Compared to freshly platelets, more than 50% platelets stored at RT for 24 hours were cleared from circulation within 5 min after transfusion (**Fig. 1C**). Platelets stored at RT for two days were cleared completely within 5 minutes after transfusion, suggesting that RT storage leads to rapid clearance, presumably as a result of previously documented “storage lesions” ^24, 25^. Consistent with previous findings ^26^, platelets stored at RT for two days or longer became completely unresponsive to stimulation with the thrombin receptor agonist PAR4 peptide (**Fig. S1A**), and lost normal integrity, because platelet volume doubled after being stored at RT for two days (**Fig. S1B**).

**Figure 1.**
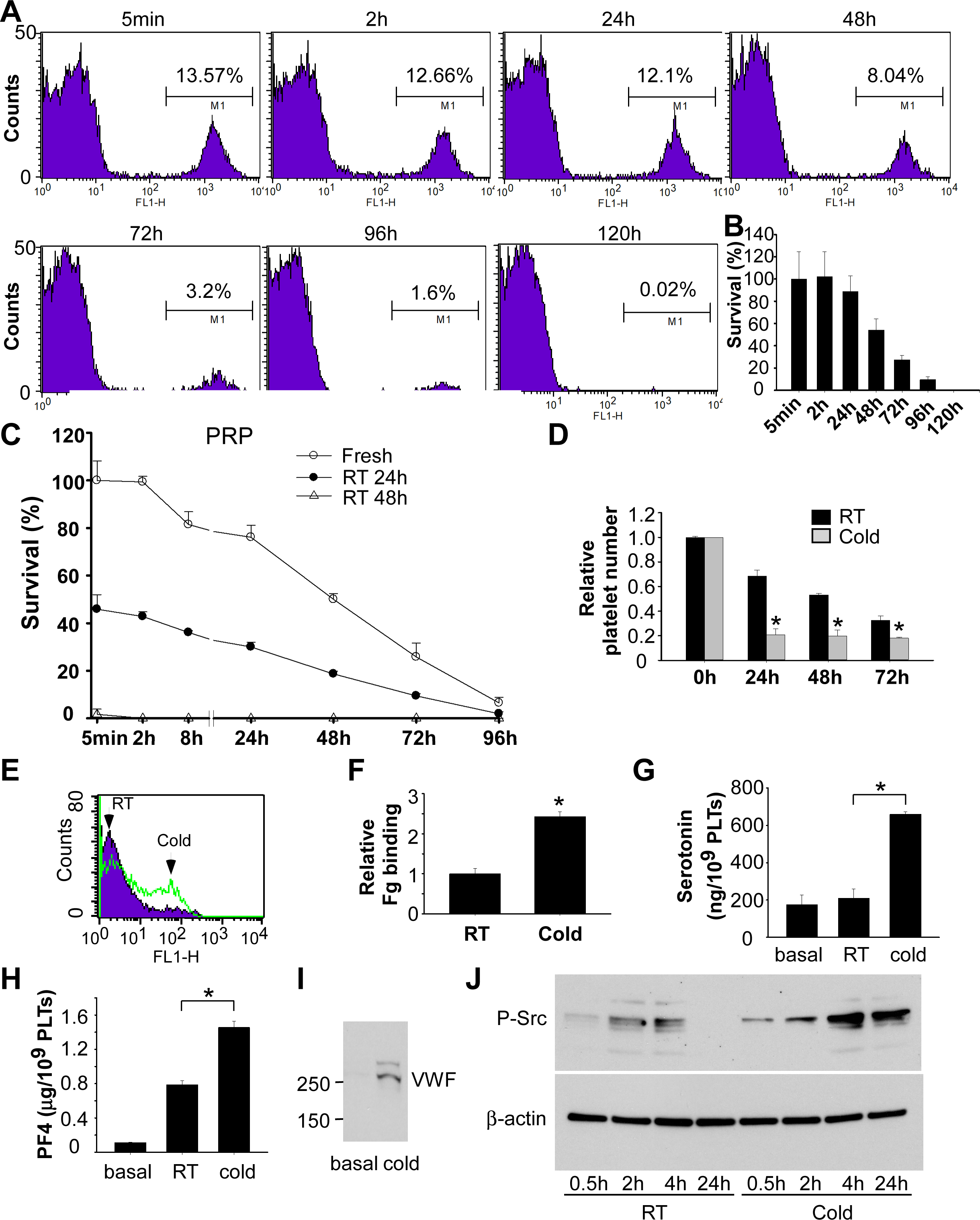
RT storage caused lesions to platelets and cold storage induced platelet activation. **A and B**, Establishment of an assay detecting life span of transfused platelets. C57BL/6J mice were injected retro-obitally with washed platelets (2.5 × 10^8^ per mouse) from C57BL/6-Tg(CAGEGFP)1Osb/J mice. Blood (<60 μl each time) was collected from mice and platelets were analyzed by flow cytometry. Quantitative results were expressed as percentage of survived transfused platelets (**B**) (percentage of GFP positive cells/percentage of GFP positive cells at 5 min after transfusion; mean ± SD; n=5). **C**, PRP from C57BL/6J mice was stored at RT for 24 h or 48 h. PRP was warmed in 37°C for 15 min and injected retro-orbitally injected to 8 weeks old C57BL/6J mice (2.5 × 10^8^ in 0.2 ml per mouse). Blood samples were collected at various time points after transfusion. Platelets were isolated by centrifugation and analyzed by flow cytometry. Survival of the transfused platelets was normalized to the survival of transfused fresh platelets at 5 min after transfusion (n=4∼6). **D**, Washed platelets (3 × 10^8^/ml) from C57BL/6J mice were stored at RT or 4°C for 24h. Platelet counts were measured with a HEMAVET HV950FS multispecies hematology analyzer at indicated time and shown as relative platelet numbers to the basal platelet count. **E** and **F**, Washed platelets from C57BL/6J mice were stored at RT or 4°C for 24 h. Integrin αIIbβ3 activation was analyzed by FITC-labeled fibrinogen binding using flow cytometry. A representative flow cytometry plots (**E**) and quantification of fibrinogen (Fg) binding relative to RT (**F**) were shown. **G and H**, Washed platelets from C57BL/6J mice were stored at RT or 4°C for 24 h. The amount of serotonin (**G**) and PF4 (**H**) in supernatant were measured as described in the methods. **I**, Washed platelets from C57BL/6J mice were stored at 4°C for 24 h. Platelets were centrifuged and VWF in supernatant was detected by Western blot with a rabbit polyclonal antibody. **J**, Washed platelets from C57BL/6J mice were stored at RT or 4°C for indicated time and solubilized with SDS-PAGE sample buffer. Phosphorylation of Src was detected by Western blot with an antibody recognizing phosphorylated Src. An antibody against β-actin was used to verify equal loading. Shown is a representative figure of three independent experiments.

### Cold storage resulted in platelet activation and aggregation

Cold-stored platelets have a better hemostatic function than RT-stored platelets. Unfortunately, cold-stored platelets are cleared rapidly after transfusion. To develop a method preventing the rapid clearance of cold-stored platelets, we first need to understand the mechanism by which cold storage leads to rapid clearance. We found that when washed platelets were incubated at 4°C for over 24 hours, small aggregates were visible in the platelet suspension (**Fig. S2A**). Single platelet count confirmed that platelets were aggregated during cold storage using a HEMAVET HV950FS hematology analyzer. Platelet counts reduced significantly after storage at 4°C for 24h or longer (**Fig. 1D**). These data suggest that platelet aggregation occurred during cold storage.

Platelet aggregation is a consequence of integrin activation. Thus, we measured fibrinogen binding to platelets to determine whether cold storage could induce integrin activation. As expected, fibrinogen binding to cold-stored mouse platelets increased compared to RT-stored platelets (**Fig. 1E** and **1F**). Thus, cold storage induces integrin activation in mouse platelets.

### Cold storage induced platelet secretion and Src kinase phosphorylation

Platelet activation is inevitably accompanied by secretion. Thus, we investigated whether cold storage could elicit platelet secretion. Secretion from dense granules was evaluated by measuring serotonin in supernatant and secretion from α granules was determined by measuring platelet factor 4 (PF4) and von Willebrand factor (VWF) in supernatant from cold-stored platelets. Indeed, cold storage elicited secretion from both dense (**Fig. 1G**) and α granules (**Fig. 1H** and **1I**).

Platelet activation involves multiple signaling pathways including the PI3K/Akt pathway, MAP kinase pathway, and Src family kinase pathway, leading to phosphorylation of these kinases. We found that Src phosphorylation dramatically increased after cold storage (**Fig. 1J**).

### RGDS treatment inhibited platelet activation by cold storage and reduced rapid clearance after cold storage

RGDS peptide, an integrin inhibitor, inhibited cold storage-induced platelet aggregation (**Fig. S2B**) and reduction in platelet counts (**Fig. 2A**). Consistent with previous findings, cold storage resulted in rapid clearance of transfused platelets (**Fig. 2B**). More than 80% of transfused platelets were cleared within 5 min after injection. To determine whether platelet activation contributes to rapid clearance after cold storage, washed platelets were pre-treated with RGDS peptide and then stored at 4°C for 24 hours. RGDS peptide largely prevented rapid clearance of cold-stored platelets. Platelets stayed in circulation for at least 2 hours after transfusion (**Fig. 2B**). These data suggest that platelet activation and aggregation contribute to cold storage-induced rapid clearance. Platelets in PRP had a better outcome of clearance than washed platelets after cold storage. Addition of RGDS peptide also improved platelet clearance in PRP after cold storage (**Fig. 2C**)

**Figure 2.**
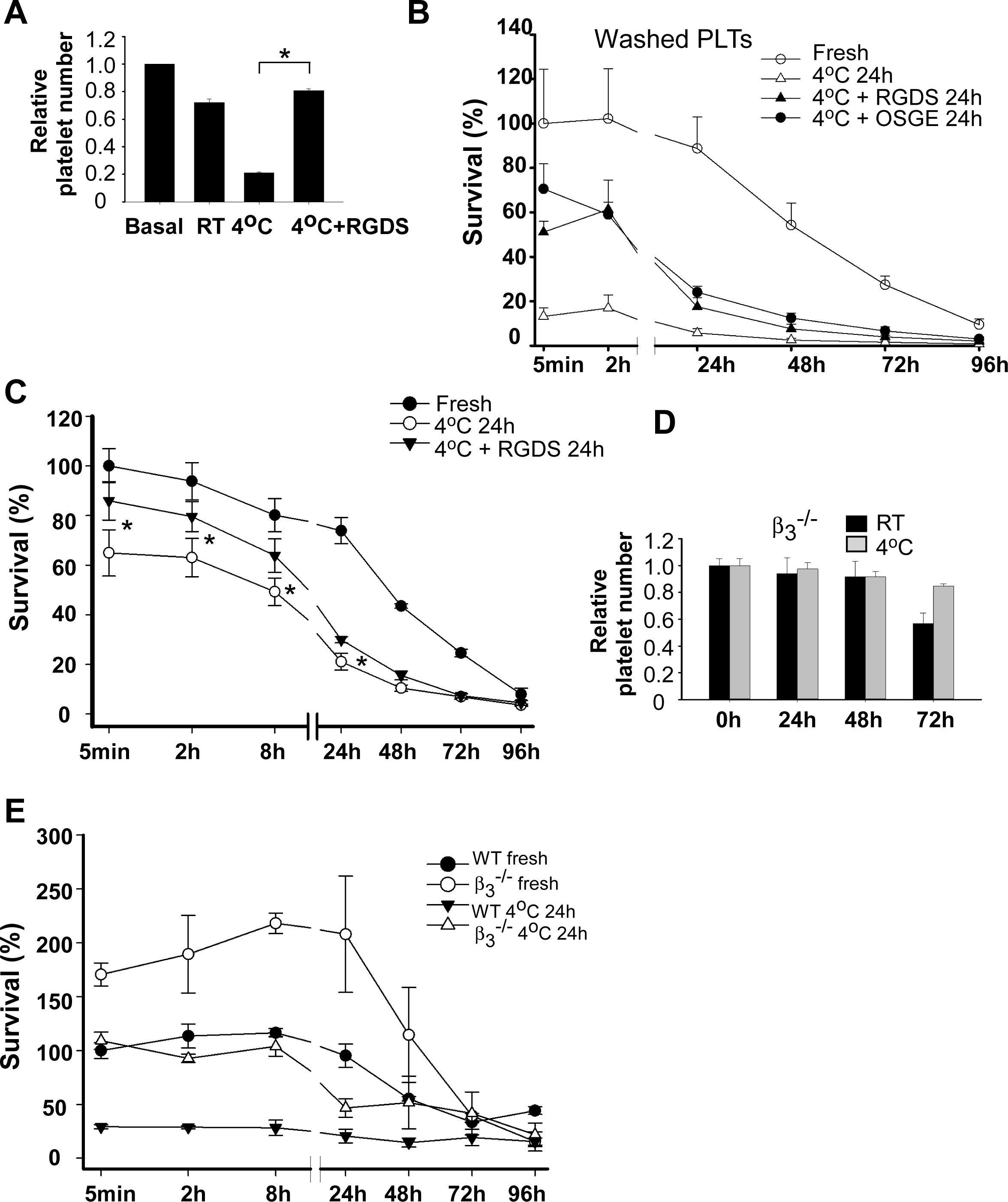
Cold storage-induced platelet activation and clearance required integrin activation, which involved the GPIb-IX signaling. **A**, Washed platelets (3 × 10^8^/ml) from C57BL/6J mice were preincubated with vehicle or RGDS (1mM) for 15 min at RT and then stored at RT or 4°C for 24 h. Platelet counts were measured with a HEMAVET HV950FS multispecies hematology analyzer at indicated time and shown as relative platelet numbers to the basal platelet count. The statistical differences were examined by Student t test. Values are means ± s.d. (n=3). *, p<0.05. **B**, Washed platelets from the C57BL/6-Tg(CAGEGFP)1Osb/J mice were preincubated with vehicle, RGDS (1 mM), or O-sialoglycoprotein endopeptidase (OSGE) for 15 min at RT and then stored at 4°C for 24 h. Platelets (2.5 × 10^8^ in 0.2 ml per mouse) were pre-warmed and injected retro-orbitally into C57BL/6J mice. Blood samples were collected at various time points after transfusion. Platelets were isolated by centrifugation and analyzed by flow cytometry. Survival of the transfused platelets was normalized to the survival of transfused fresh platelets at 5 min after transfusion (n=4∼6). **C**, PRP from the C57BL/6-Tg(CAGEGFP)1Osb/J mice were preincubated with vehicle or RGDS (1 mM) for 15 min at RT and then stored at 4°C for 24 h. Platelets (2.5 × 10^8^ in 0.2 ml per mouse) were pre-warmed and injected retro-orbitally into C57BL/6J mice. Blood samples were collected at various time points after transfusion (n=4). **D**, Washed platelets (3 × 10^8^/ml) from integrin β3^-/-^ mice were stored at RT or 4°C for indicated time and platelet counts were measured. **E**, Washed platelets from the β3 deficient mice and WT controls were stored at 4°C for 24 h and then injected retro-orbitally into C57BL/6-Tg(CAGEGFP)1Osb/J mice. Survival of the transfused platelets was normalized to the survival of transfused fresh platelets at 5 min after transfusion (n=4)

### Platelets from integrin β3 deficient mice were protected from cold storage induced rapid clearance after transfusion

Next, we used integrin β3 deficient mice to determine the role of integrin in cold storage-elicited platelet activation and clearance. Cold storage failed to induce platelet aggregation and reduction in platelet counts in the β3 deficient mice (**Fig. 2D**). To determine the role of β3 deficiency in cold storage-induced rapid clearance, washed platelets from the β3 deficient or WT mice were stored at 4°C for 24 hours and then transfused into the C57BL/6-Tg(CAGEGFP)1Osb/J mice. Rapid clearance by cold storage was largely inhibited in β3 deficient platelets (**Fig. 2E**). These data demonstrate that rapid clearance of platelets after cold storage mainly depends on integrin activation. However, β3 deficient platelets were cleared faster than fresh platelets after 24 hours of transfusion suggesting that other mechanisms may be involved. Interestingly, mice transfused with fresh β3 deficient platelets had much higher concentrations of transfused platelets in circulation than the mice transfused with wild type platelets. This might be due to reduced adhesion of the β3 deficient platelets to vasculature after transfusion.

### Cold-induced platelet activation was independent of the ADP and TXA2 dependent pathways

Next, we explore the mechanism of cold storage-elicited platelet activation. Platelet activation involves a series of rapid positive feedback loops that amplify initial activation signals and enable robust platelet recruitment and thrombus growth. Release of ADP and TXA2 are two major positive feedback mechanisms of platelet activation. To determine whether cold storage-induced platelet activation involves these two feedback mechanisms, we examined whether platelet activation by cold storage was inhibited in platelets from P2Y12 and TP deficient mice. Similar to wild type platelets, P2Y12^-/-^ and TP^-/-^ platelets aggregated when stored at 4°C for 24h (**Fig. 3A** and **3B**). Accordingly, platelet counts were reduced significantly. These data indicate that ADP and TXA2 are not required for cold temperature-induced platelet activation. Because cold storage induced platelet secretion, we examined whether secretion is involved in cold temperature-elicited platelet activation and aggregation. Muc13-4 plays a critical role in mediating platelet secretion. However, low temperature-induced platelet aggregation was not affected in the Unc13d(Jinx) mice that were deficient in Munc13-4 (**Fig. 3C**). Reduction in platelet counts was partially reversed in Gq^-/-^ platelets (**Fig. 3D**), suggesting that Gq signaling is involved in cold-induced platelet activation.

**Figure 3.**
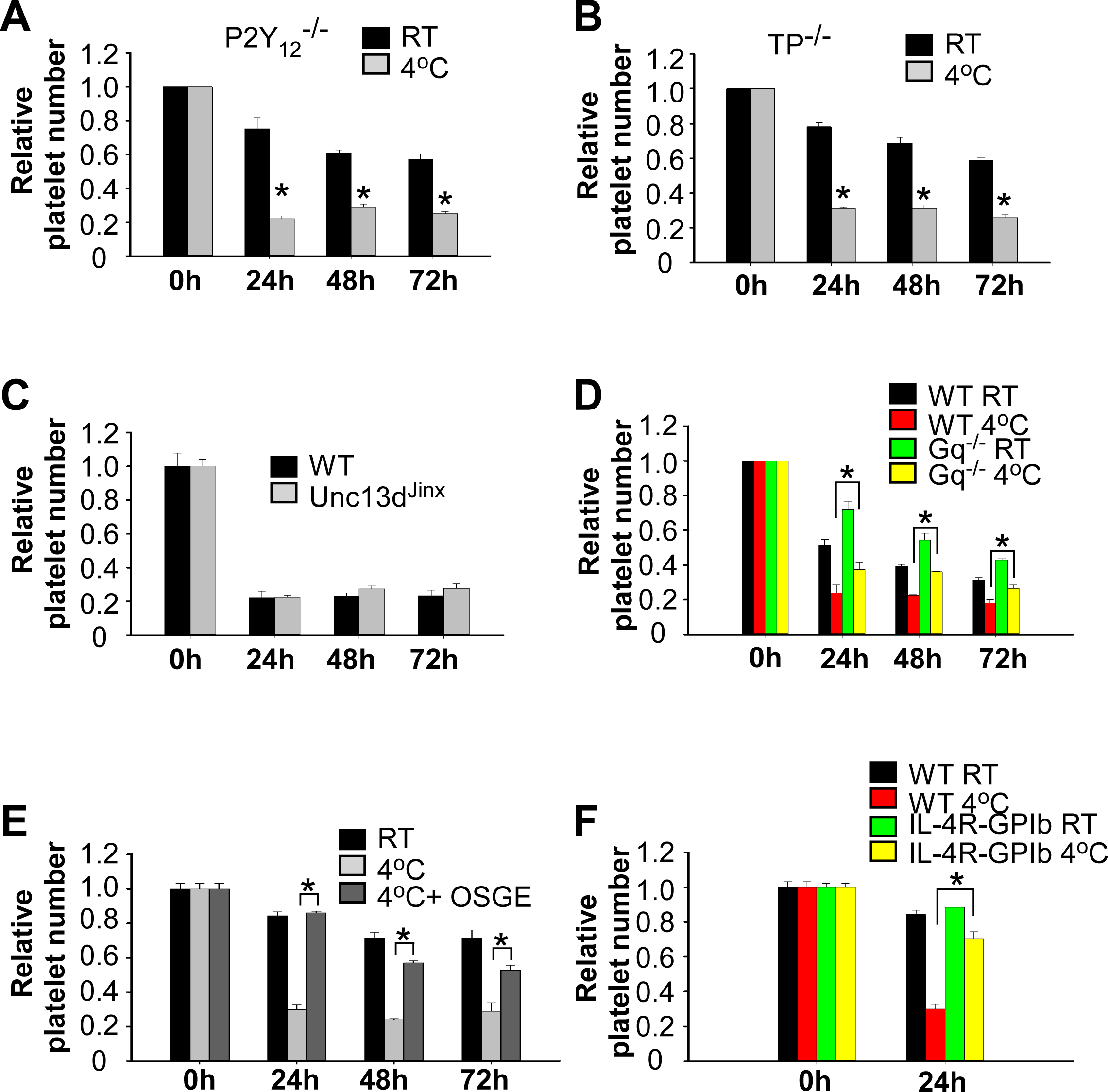
Cold storage-induced platelet activation involved GPIb-IX signaling. **A** and **B**, Washed platelets (3 × 10^8^/ml) from P2Y12 (**A**) or TP (**B**) deficient mice were stored at RT or 4°C for indicated time. Platelet counts were measured and shown as relative platelet number to the platelet count at 0 h. **C**, Washed platelets (3 × 10^8^/ml) from Munc13-4 deficient mice or wild-type controls were stored at 4°C for indicated time. Platelet counts were measured and shown as relative platelet number to the platelet count at 0 h. **D**, Washed platelets (3 × 10^8^/ml) from Gq deficient mice or littermate wild-type controls were stored at RT or 4°C for indicated time and single platelet counts were measured. **E**, Washed platelets were pre-treated with or without sialoglycoprotein endopeptidase (OSGE) for 15 min at RT and then stored at 4°C or RT. **F**, Washed platelets (3 × 10^8^/ml) from IL4Rα/GPIbα mice or wild-type controls were stored at RT or 4°C for 24 h. Single platelet counts were measured and shown as relative platelet number to the platelet count at 0 h. The statistical differences were examined by Student t test. Values are means ± s.d. (n=3). * p<0.05.

### GPIb signaling played an important role in cold storage-induced platelet activation

Previous studies have shown that the glycoprotein Ibα plays a critical role in cold storage-induced rapid clearance of transfused platelets ^17^. We used different approaches to evaluate the role of GPIb signaling in cold storage-elicited platelet activation. Pretreatment of platelets with an O-sialoglycoprotein endopeptidase (OSGE), which strips the extracellular domain of GPIbα ^27^, prevented cold storage-induced platelet aggregation (**Fig. 3E**). Similarly, cold storage-induced platelet aggregation was reduced in GPIbα deficient mice expressing a chimeric human IL-4 receptorα/GPIbα (IL4Rα/GPIbα) (**Fig. 3F**). Our findings are consistent with a role for GPIbα signaling in cold storage-elicited rapid clearance ^17^.

### EGTA prevented cold storage-induced platelet activation

Although integrin inhibitors efficiently prevented rapid clearance after cold storage, they could not be developed as reagents for cold storage of platelets. An effective method for platelet cold storage must meet two prerequisites: i) Platelets must remain active and ii) the agents must not compromise patient safety. Therefore, the desired reagents that allow platelet cold storage should inhibit cold storage-induced platelet activation, but this inhibition must be reversible. In searching for a method that fits all the prerequisites, we unexpectedly found that a non-cell permeable calcium ion chelator, EGTA, completely inhibited cold-induced platelet aggregation and reduction in platelet counts (**Fig. 4A**). Importantly, this inhibition was reversible. When a physiological concentration of calcium ions was restored, platelets fully responded to subsequent stimulation of agonist (**Fig. 4B**).

**Figure 4.**
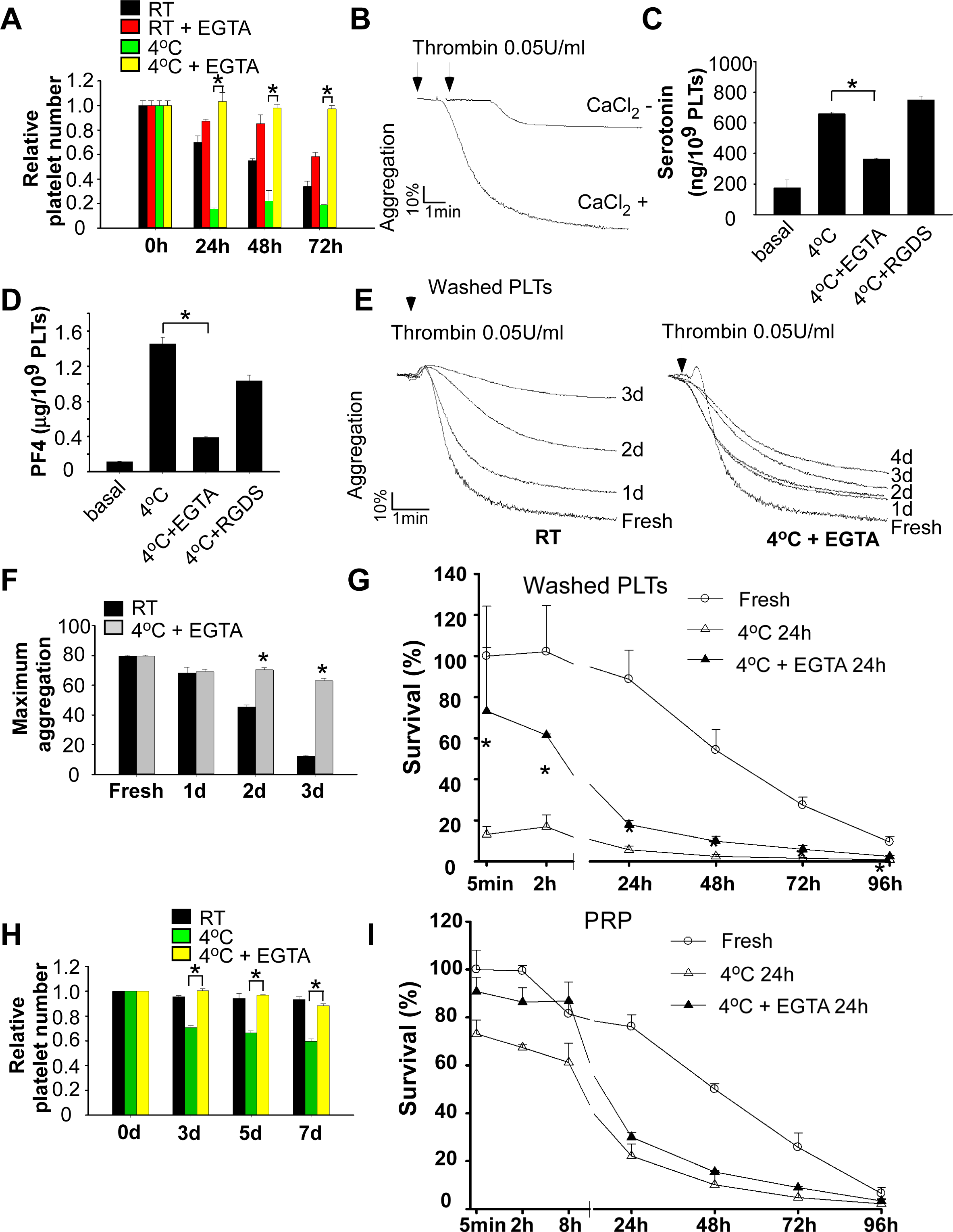
EGTA treatment protected against cold-induced platelet activation and rapid clearance. **A**, Washed platelets (3 × 10^8^/ml) from C57BL/6J mice were preincubated with vehicle or EGTA (100 µM) for 15 min at RT and then stored at RT or 4°C for indicated time. Platelets accounts were measured and shown as relative to the counts at 0 h. The statistical differences were examined by Student t test. Values are means ± s.d. (n=3). * p<0.05. **B**, Washed platelets from C57BL/6J mice were preincubated with EGTA for 15 min at RT and then stored 4°C for 24 h. Platelets were warmed in 37°C for 15 min and added with 1 mM CaCl_2_. Platelet aggregation was induced by adding 0.05 U/ml thrombin. **C** and **D**, Washed platelets from C57BL/6J mice were pretreated with 100 µM EGTA or 1 mM RGDS for 15 min. Platelets were then stored at 4°C for 24 h. The amount of serotonin (**C**) and PF4 (**D**) in supernatant were measured as described in the methods. **E** and **F**, Washed platelets from C57BL/6J mice were stored at RT for indicated lengths of time. Platelets were also stored at 4°C in the presence of EGTA (100 μM). Platelets were prewarmed at 37°C for 15 min and added with 1 mM CaCl_2_. Aggregation was induced by addition of thrombin. **G**, Washed platelets from C57BL/6-Tg(CAGEGFP)1Osb/J mice were preincubated with EGTA for 15 min at RT and then stored at cold for 24 h. Pre-warmed platelets (2.5 × 10^8^ in 0.2 ml per mouse) were injected retro-orbitally injected to 8 weeks old C57BL/6J mice. Blood samples were collected at various time points after transfusion (n=4-6). **H**, PRP from C57BL/6J mice were stored at RT or 4°C in the presence of absence of EGTA for 24 h, 48 h, and 72 h. Platelet counts were measured with a HEMAVET HV950FS multispecies hematology analyzer at indicated time and shown as relative platelet numbers to the basal platelet count. **I**, PRP from GFP mice were preincubated with vehicle or EGTA for 15 min at RT and then stored at 4°C for 24 h. Pre-warmed PRP (2.5 × 10^8^ platelets in 0.2 ml per mouse) was injected retro-orbitally to 8 weeks old C57BL/6J mice (n=3-6).

EGTA inhibited platelet secretion elicited by cold storage (**Fig. 4C** and **4D**). In contrast, although RGDS peptide inhibited cold-induced platelet aggregation, it did not inhibit platelet dense secretion by cold storage. These data indicate that EGTA inhibits cold storage-induced platelet secretion through an integrin-independent mechanism. These data also suggest that Ca^2+^ binding to the extracellular membrane of platelets is not only required for integrin ligand binding function, but also elicits an early signaling event leading to integrin activation and platelet secretion.

### EGTA treatment inhibited rapid clearance of cold-stored platelets after transfusion

Platelets stored at 4°C with EGTA maintained reactivity to agonists (**Fig. 4E** and **4F**). Treatment of washed platelets with EGTA significantly inhibited rapid clearance after cold storage for 24 hours (**Fig. 4G**). Similar to RGDS, EGTA did not dramatically prevent platelet clearance after 24 hours of transfusion.

### EGTA inhibited cold-elicited platelet aggregation in PRP and rapid clearance

Cold storage also resulted in platelet aggregation and reduction in platelet counts in PRP, although at a much less extent compared to washed platelets (**Fig. 4H**). These data are consistent with previous findings that clots were often observed in the cold-stored platelet products, which led to substantial wastage ^28^. Addition of EGTA inhibited reduction in platelet counts during cold storage (**Fig. 4H**). EGTA treatment also reduced platelet clearance after cold storage (**Fig. 4I**).

### Apoptosis of cold stored platelets contributed to clearance

It is known that storage of platelets at RT induces apoptosis. To determine whether apoptosis is also involved in cold storage-induced rapid clearance, we determined whether cold storage causes apoptosis of platelets. Caspase-3 is activated in the apoptotic cell both by extrinsic (death ligand) and intrinsic (mitochondrial) pathways ^29, 30^. Thus, we measured caspase-3 cleavage to determine whether apoptosis occurred in cold stored platelets. Clearly, cold storage caused caspase-3 cleavage in mouse platelets (**Fig. 5A**). Cold storage-elicited caspase-3 cleavage is abolished by pre-incubation of platelets with RGDS or EGTA (**Fig. 5A**), as well as integrin β3 deficiency (**Fig. 5B**). These data suggested that cold storage indeed induced platelet apoptosis, which required integrin activation. To determine whether apoptosis contributes to cold storage-induced platelet clearance, PRP was prepared from caspase-3 deficient mice or WT controls and stored at 4°C for 24 hours. There is no difference between fresh caspase-3 deficient platelets and WT controls. However, the number of the stored caspase-3 deficient platelets in circulation was significantly increased at 24, 48, and 72 hours post transfusion compared to WT controls (**Fig. 5C**).

**Figure 5.**
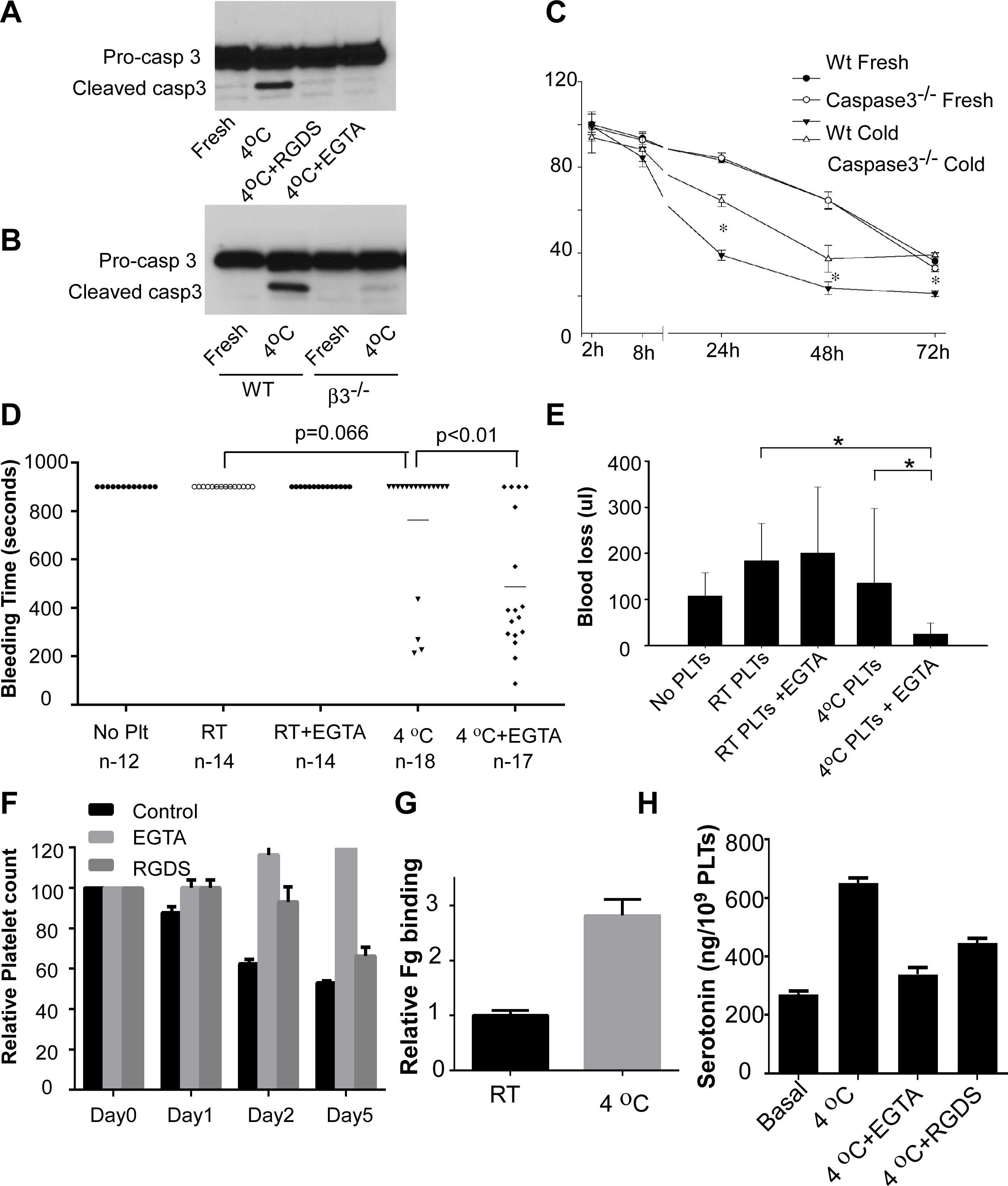
Apoptosis contributed to cold storage-induced clearance and cold-stored platelets protected against bleeding in GPIb α deficient mice. **A**, Washed platelets from C57BL/6J mice were preincubated with vehicle, RGDS (1 mM) or EGTA (0.1 mM) for 5 min and then stored at 4°C for 4 hours and solubilized with SDS-PAGE sample buffer. Pro-caspase-3 and cleaved caspase-3 were detected by Western blot. **B**, Washed platelets from β3 deficient mice or WT controls were stored at 4°C for 4 hours and solubilized with SDS-PAGE sample buffer. Pro-caspase-3 and cleaved caspase-3 were detected by Western blot. **C**, PRP from caspase-3 deficient mice and WT controls were stored at 4°C for 24 h and then injected retro-orbitally into C57BL/6-Tg(CAGEGFP)1Osb/J mice. Survival of the transfused platelets was normalized to the survival of transfused fresh WT platelets at 2h after transfusion (n=4, * p<0.01). **D** and **E**, 7∼8 weeks old mice deficient in GPIbα received RT-stored platelets or cold-stored platelets in the presence or absence of EGTA (1 × 10^9^ platelets in 0.2 ml PRP per mouse by retro-orbital injection). Mouse tail-bleeding times (**D**) and blood loss (**E**) were determined as described in the method. **F**, PRP from a healthy donor was preincubated with vehicle or EGTA (2 mM) or RGDS (1 mM) for 15 min at RT and then stored at RT or 4°C for indicated time. Platelets accounts were measured and shown as relative to the counts at 0 h. **G**, Washed platelets from a healthy donor was stored at RT or 4°C for 24 h. Integrin αIIbβ3 activation was analyzed by FITC-labeled fibrinogen binding using flow cytometry. Quantification of fibrinogen (Fg) binding relative to RT were shown. **H**, Washed human platelets were pretreated with EGTA or RGDS for 15 min. Platelets were then stored at 4°C for 24 h. The amount of serotonin in supernatant were measured as described in the methods.

### EGTA-treated, cold stored platelets efficiently prevented bleeding in GPIbα deficient mice

To determine whether EGTA-treated, cold-stored platelets can efficiently prevent bleeding, EGTA-treated PRP was stored at 4°C for 48 hours and then transfused into GPIbα deficient mice. GPIb-IX complex is essential for platelet production and hemostasis. GPIbα deficiency or loss of function leads to Bernard-Soulier syndrome (BSS) ^31^. Similar to BSS patients, mice deficient in GPIbα have thrombocytopenia, giant platelets, and bleeding tendency. Injection of EGTA-treated, cold-stored platelets in PRP into GPIbα deficient mice significantly shortened tail-bleeding times (**Fig. 5D**) and reduced blood loss (**Fig. 5E**). EGTA-treated, cold-stored platelets had a better outcome than the cold-stored platelets in the absence of EGTA. In contrast, injection of the same amount of RT-stored platelets, with or without EGTA, had no effects in preventing blood loss and reducing tail-bleeding times in the GPIbα deficient mice.

### Platelet counts were reduced in human PRP after cold storage

To determine whether cold storage causes aggregation in human platelets, washed platelets or PRP prepared from healthy donors using ACD as anticoagulant was stored at 4°C, in the absence or presence of EGTA or RGDS peptide. Platelet counts were monitored at various time points. Platelet counts decreased after storage at 4°C in a time-dependent manner, which was inhibited by EGTA and RGDS, in both PRP (**Fig. 5F**) or washed platelets (**Fig. S3)**. Fibrinogen binding to platelets increased after cold storage (**Fig. 5G** and **Fig. S4**). Cold storage also induced platelet secretion (**Fig. 5H**). These data suggest that similar to mouse platelets, human platelets are activated during cold storage.

### EGTA inhibited cold-elicited human platelet aggregation and better preserved platelet function

To determine whether cold-stored platelets in the presence of EGTA preserve the hemostatic function better than the RT-stored platelets, we compared platelet response to ADP and PAR1 peptide between RT-stored platelets and cold-stored platelets. RT-stored platelets lost response to ADP after 3 days (**Fig. 6A** and **6B**). In contrast, the EGTA-treated, cold-stored platelets maintained response to ADP even after 9 days. EGTA-treated, cold-stored platelets also had a better response to PAR1 peptide than the RT-stored platelets (**Fig. 6C** and **6D**).

**Figure 6.**
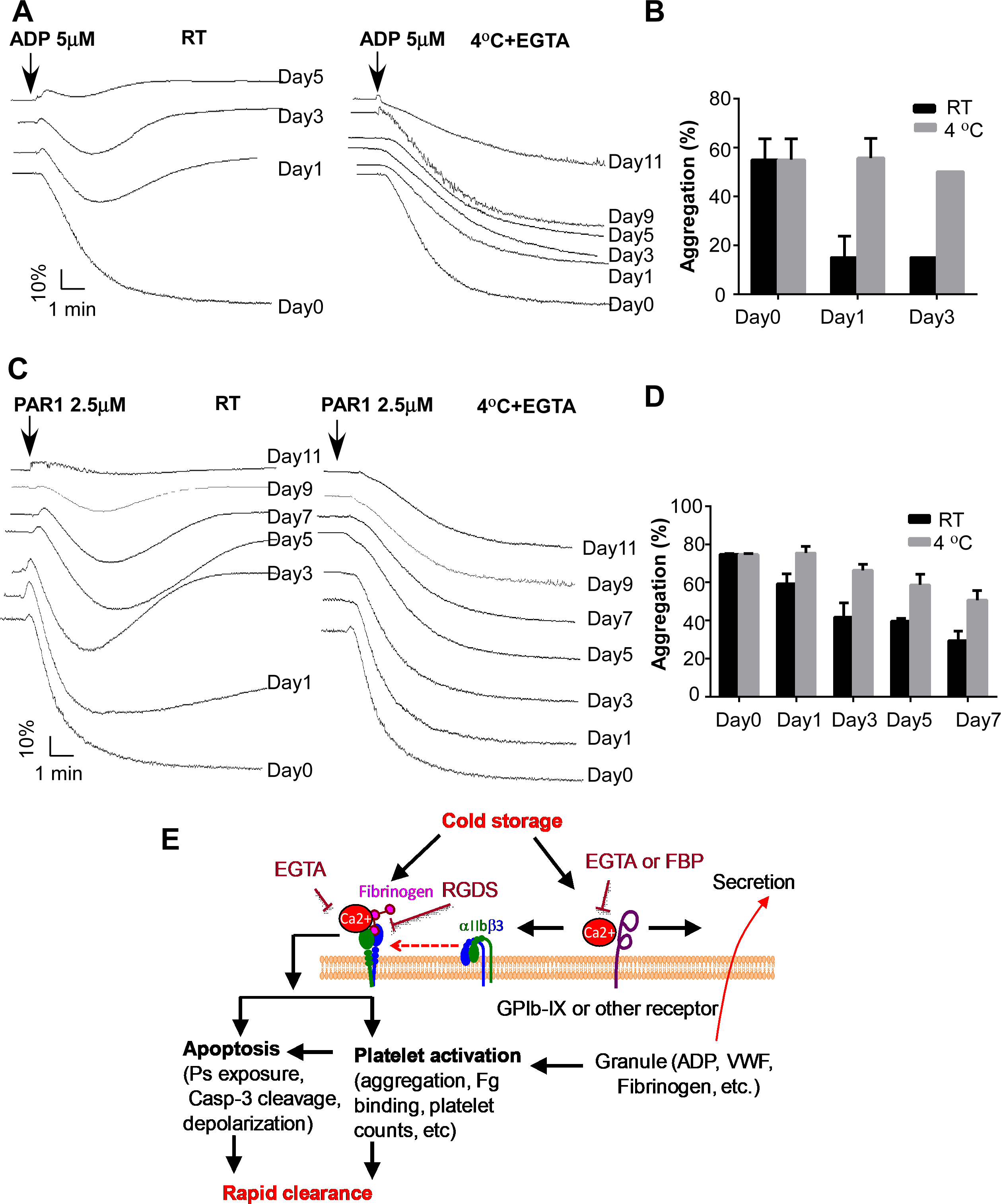
Cold-stored human platelets preserve better function than RT-stored platelets. **A** and **B**, PRP from healthy donors was added with buffer or EGTA (2 mM) and incubated at RT for 15 min, and then stored at RT or 4°C for indicated lengths of time. Platelets were prewarmed at 37°C for 15 min and added with 1 mM CaCl_2_. Aggregation was induced by addition of 5 μM ADP (n=3). **C** and **D**, PRP from healthy donors was added with buffer or EGTA (2 mM) and incubated at RT for 15 min, and then stored at RT or 4°C for indicated lengths of time. Platelets were prewarmed at 37°C for 15 min and added with 1 mM CaCl_2_. Aggregation was induced by addition of 2.5 μM PAR1 peptide (n=3). **E**, A schematic of cold storage induced platelet clearance.

## Discussion

It is known that refrigerated platelets are cleared rapidly after transfusion ^2-4, 17-20^. The mechanism by which cold storage induces platelet rapid clearance is not fully understood. Our data suggest that rapid clearance of cold-stored platelets is largely due to integrin activation. We demonstrate that cold storage elicits platelet activation, which requires binding of calcium ion to the extracellular membrane of platelets. Incubation of platelets with a cell impermeable calcium ion chelator EGTA inhibited cold-induced platelet activation and aggregation and prevented rapid clearance of cold-stored platelets (**Fig. 6E)**.

Previous studies have shown that incubation of platelets at cold temperature induces morphological changes ^12, 13, 32^ and some degree of activation, including increase in intracellular Ca^2+^ concentration ^14^, cytoskeletal rearrangements ^15^, and spontaneous aggregation after they are warmed and stirred at 37°C immediately after removal from cold environment ^16^. In this study, we provide further evidence that cooling induces integrin activation. Cold storage gradually enhanced fibrinogen binding to platelets. Single platelet counts reduced dramatically after platelets were stored at 4°C for over 24 hours and small aggregates were visible in suspension. The aggregates formed during cold storage are different from the aggregation induced by agonists. Most aggregates formed during cold storage could be resuspended after prewarming the platelets at 37°C and pipetting. However, resuspension did not prevent clearance after cold storage. We conclude that cold storage-induced rapid clearance is largely due to integrin activation, because deficiency of integrin β3 dramatically restored circulation of cold-stored platelets and an integrin inhibitor RGDS peptide significantly inhibited cold storage-induced rapid clearance. Our data suggest that apoptosis is one of the major mechanism that contributes to platelet clearance. Apoptosis resulting from cold storage appears to be downstream from integrin activation, since cold storage-elicited cleavage of caspase-3 was abolished by the integrin inhibitor RGDS peptide and β3 deficiency.

Previous studies suggest an important role of GPIbα in low temperature-induced platelet clearance. Consistent with previous findings, we found that treatment of platelets with O-sialoglycoprotein endopeptidase, which strips the extracellular domain of GPIbα ^27^, or deficiency of GPIbα, significantly inhibited rapid clearance of platelets after cold storage (**Fig. 2B**). Because cold storage-induced clearance is largely αIIbβ3 dependent, we speculate that the GPIb-IX signaling plays an important role in cold storage-induced platelet activation and that contribution of GPIbα to rapid clearance is due to GPIb-IX signaling-mediated αIIbβ3 activation. This conclusion was supported by the data that (1) treatment of platelets with O-sialoglycoprotein endopeptidase inhibited cold-induced platelet aggregation and reduction in platelet counts; (2) platelets from the IL4Rα/GPIbα deficient mice, which express a chimeric human IL-4 receptorα/GPIbα, were resistant to cold storage-induced platelet aggregation; (3) cold storage-induced fibrinogen binding to platelets was inhibited in platelets from the IL4Rα/GPIbα mice (**Fig. S5)**. This conclusion is also supported by previous studies that VWF binding to platelets was induced during cold storage and that refrigeration-induced binding of VWF to platelets facilitates their rapid clearance by inducing GPIbα-mediated signaling ^20, 33, 34^. GPIb-IX complex is a sensor for shear stress-induced platelet activation ^35^. Shear stress induces unfolding of a mechanosensory domain in GPIbα, leading to intracellular signaling and rapid platelet clearance ^36^. Low temperature might cause stress to the cells similar as shear stress, resulting in unfolding of the GPIbα mechanosensory domain on the platelet, which leads to integrin activation and clearance. However, cold storage-induced platelet activation was not completely GPIbα dependent, because lack of GPIbα or VWF, and pretreatment of platelets with O-sialoglycoprotein endopeptidase only partially reversed rapid clearance after cold storage.

Since cold storage-induced aggregation was partially inhibited in the Gq deficient platelets, we hypothesized that Ca^2+^ mobilization plays a role in cold storage-induced platelet dysfunction. We tested this hypothesis using a cell-permeable calcium ion chelator BAPTA-AM. A cell impermeable chelator EGTA was used as a control for this experiment. Surprisingly, EGTA completely inhibited storage-induced aggregation and reduction in platelet counts. Since platelets were suspended in a modified Tyrodes’ solution in the absence of Ca^2+^, these data suggest that Ca^2+^ binding to the extracellular membrane of platelets is required for the cold storage-induced platelet activation. Earlier studies have shown that treatment of platelets with calcium ion chelators such as EDTA irreversibly dissociate αIIbβ3 complex, leading to irreversible loss of platelet aggregation function ^37, 38^. Dissociation of αIIbβ3 complex by EDTA requires platelet incubation with EDTA at 37°C. While incubation of platelets with EGTA at room temperature inhibited agonist-induced fibrinogen binding to platelets (**Fig. S6A**), this process is reversible. Addition of CaCl_2_ completely reversed fibrinogen binding to platelets (**Fig. S6B**). Similarly, addition of CaCl_2_ completely restored aggregation of EGTA-treated platelets in both mice and humans (**Fig. S7**). Platelet spreading on fibrinogen was also inhibited by EGTA and addition of CaCl_2_ completely reversed platelet spreading (**Fig. S8A** and **S8B**). Therefore, incubation of platelets with EGTA under our conditions did not cause irreversibly dissociate αIIbβ3 complex. Cold storage-elicited platelet secretion was not inhibited by the RGDS peptide, suggesting that cold-induced platelet secretion is integrin independent. These data are consistent with our previous findings that agonist-induced platelet secretion is integrin independent ^39^. Interestingly, cold storage-induced platelet secretion was inhibited by EGTA. Thus, Ca^2+^ binding to platelets not only regulates integrin function, but also affects an early signaling event, which leads to platelet secretion and integrin activation.

Since 1960s, platelet products for transfusion are routinely stored in di-(2-ethylhexyl) phthalate (DEHP) plasticized polyvinyl chloride bags at 22°C to 24°C under continuous gentle agitation ^40, 41^. Human platelets stored at RT for three days lose responsiveness and are less effective in preventing bleeding in thrombocytopenic patients ^3^. Platelets stored at RT undergo a series of changes in morphology and function, leading to platelet storage lesions, including loss of the discoidal shape, release of granule contents, PS exposure, and modifications of glycoprotein patterns on the surface ^41^. Consistent with observations from human platelets, we show that mouse platelets lost response to agonist stimulation and were cleared immediately after they were stored at RT for two days. Because of the problems caused by RT storage, in 2015, the FDA and the American Association of Blood Banks (AABB) approved a procedure to use 3-day cold-stored platelets without agitation and bacterial testing for patients with actively bleeding trauma because of the advantages of cold-stored platelets over RT-stored platelets ^28^. Unfortunately, the approved cold storage method resulted in an 80.9% disposal rate of platelet products. One major reason causing disposal of cold-stored platelets is the formation of clots ^28^. Our data suggest that platelet aggregation during cold storage is at least one of the major reasons causing the formation of “clots”. We show that platelets are aggregated and platelet counts dropped in the PRP prepared using the same procedure for blood transfusion in both mouse and human platelets.

Administration of a glycoprotein αIIbβ3 antagonist Abciximab, a Fab fragment of the chimeric human-murine monoclonal antibody 7E3, induces thrombocytopenia in some patients. Thrombocytopenia can occur in up to 12% of patients after repeat exposure and 1-2% of patients on first exposure due to the presence of naturally-occurring antibodies that recognize the murine elements of the drug ^42^. It has been reported that thrombocytopenia induced by Abciximab occurs in 2-4 hours after exposure to Abciximab ^43^. Severe thrombocytopenia induced by Abciximab appears to be more common when re-exposure occurs within 2 weeks ^42, 43^. To determine whether EGTA-treated platelets could cause thrombocytopenia through alloimmunity, C57BL/6J mice were transfused with EGTA-treated platelets twice in two weeks. Platelet counts were monitored at 4 hours and 24 hours after the second transfusion of EGTA-treated platelets. The mice transfused with EGTA-treated platelets had higher platelet counts and did not develop thrombocytopenia. The mice injected with EGTA alone had similar platelet counts as controls (**Fig. S9**).

It has been reported previously that platelets isolated from blood using EDTA as an anticoagulant lost discoid shape and become irregular and spherical ^44^. Few platelets prepared from blood using EDTA as an anticoagulant stay in circulation after transfusion ^44^. Platelet clearance is also much faster using EDTA as an anticoagulant than using ACD as anticoagulant. Fortunately, this effect seems to be EDTA specific, because we show that similar as RGDS peptide or endopeptidase, EGTA successfully prevented rapid clearance of cold-stored platelets. Interestingly, peripheral lymphocytes lose response to antigens if EDTA is used as an anticoagulant, but maintain normal response to antigens if EGTA is used as an anticoagulant ^45^. The underlining mechanism causing the difference between EDTA and EGTA is not clear. We show that EGTA-treated, cold-stored platelets but not the RT-stored platelets prevented bleeding in mice. If EGTA could safely prolong the circulation of cold-stored platelets in humans, it would allow cold storage of platelet products for transfusion. This method should significantly improve the shortage of platelet inventories and diminish bacterial contamination and storage lesions.

## Materials and Methods

EGTA, ADP, and CaCl_2_ were purchased from Sigma. A polyclonal antibody against phosphorylated Tyr416 of Src and a polyclonal antibody against mouse caspase-3 were from Cell Signaling Technology (Beverly, MA, USA). α-Thrombin was purchased from Enzyme Research Laboratories (South Bend, IN, USA). PAR 4 peptide AYPGKF and PAR1 peptide SFLLRN were custom-synthesized at Biomatik USA, LLC (Wilmington, DE, USA). pH SAFE Mini bag was from Blood Cell Storage, Inc (Seattle, WA).

### Mice

Mice deficient in Gα_q_ ^46^, P2Y12 ^47^, Munc13-4 ^48^, GPIbα ^49^, and integrin β3 ^46^, and GPIbα deficient mice expressing a chimeric human IL-4 receptorα/GPIbα (IL4Rα/GPIbα) ^49^ were generated as described previously. C57BL/6J mice and Tg(act-EGFP)Y01Osb mice were purchased from Jackson Laboratories. Mice were bred and maintained in the University of Kentucky Animal Care Facility following institutional and National Institutes of Health guidelines after approval by the Animal Care Committee.

### Preparation of washed murine platelets

Blood was collected from the abdominal aorta of isofluorane-anesthetized mice (8–10 weeks) using 1⁄7 volume of ACD (85 mM trisodium citrate, 83 mM dextrose, and 21 mM citric acid) as anticoagulant. The platelets were then washed once with CGS (0.12 M sodium chloride, 0.0129 M trisodium citrate, 0.03 M D-glucose, pH 6.5), resuspended in modified Tyrode’s buffer (12 mM NaHCO_3_, 138 mM NaCl, 5.5 mM glucose, 2.9 mM KCl, 2 mM MgCl_2_, 0.42 mM NaH_2_PO_4_, 10 mM HEPES, pH 7.4), and incubated for 1 h at 22 °C before use.

### Preparation of murine platelet rich plasma (PRP)

Blood was collected from the abdominal aorta of isofluorane-anesthetized mice using 1⁄7 volume of acid-citrate-dextrose (ACD-A) or 1/10 volume of 3.8% sodium citrate as anticoagulant. PRP was collected by centrifugation at 950 rpm for 10 min. PRP was further centrifuged at 2,400 rpm for 10 min to get concentrated PRP by removing some plasma.

### Preparation of human platelets

For studies using human blood, approval was obtained from the University of Kentucky institutional review board. Informed consent was provided according to the Declaration of Helsinki. Fresh blood was drawn by venipuncture from healthy volunteers, using 1/10 volume of 3.8% trisodium citrate or one-seventh of ACD-A as anticoagulant. Platelet-rich plasma (PRP) was obtained by centrifugation at 300g for 20 minutes at room temperature. To prepare washed platelets, blood was anticoagulated with ACD (1:7). PRP was further centrifuged at 1200g for 15 minutes. Platelets were then washed twice with CGS (sodium chloride 0.12 M, trisodium citrate 0.0129 M, and glucose 0.03 M, pH 6.5) containing 0.1% bovine serum albumin (BSA) and resuspended in the HEPES-buffered Tyrode solution.

### Platelets activation induced by cold storage

Washed platelets (3 × 10^8^/ml) or PRP were stored at 4°C or RT in pH SAFE Mini bags in the presence or absence of EGTA, O-sialoglycoprotein endopeptidase, or RGDS peptide. Platelet counts were monitored at various time points with a HEMAVET HV950FS multispecies haematology analyzer.

### Platelet aggregation induced by agonists after storage

Washed platelets or PRP from C57BL/6J mice (3 × 10^8^/ml) were pretreated with 100 μM EGTA or vehicle for 15 min at RT and then stored at 4°C or RT for 1 to 7 days in pH SAFE Mini bags. Platelets were then incubated at RT for 15 min or 37°C for 15 min before aggregation. Platelet aggregation measured by detecting changes in light transmission using a turbidimetric platelet aggregometer (Chrono-Log) with stirring (1000 rpm).

### Platelet transfusion

Washed platelets (1.25 × 10^9^/ml) were preincubated with vehicle, EGTA, O-sialoglycoprotein endopeptidase (OSGE), or RGDS for 15 min at RT and then stored at RT or 4°C for 24 h. Platelets were then warmed at 37°C for 15 min and retro-orbitally injected (0.2 ml/mouse) into 8 weeks old recipient mice. Blood samples (< 60 μl) were collected retro-orbitally with a heparin-containing tube at various time points after transfusion. 500 μl saline was added to the blood samples. Platelets were isolated by centrifugation at 1,000 rpm for 3 min and analyzed by flow cytometry. Ratio of GFP positive platelets in total platelets in fresh platelets group at 5 min after transfusion was set to 100% (survival 100%). Survival of the transfused platelets was normalized to the survival of transfused fresh platelets at 5 min after transfusion.

For PRP transfusion experiments, concentrated PRP (1.25 × 10^9^/ml) were preincubated with vehicle, EGTA, or RGDS for 15 min at RT and then stored at RT or 4°C for 24 h or 48 h. The PRP was warmed in 37°C for 15 min and was retro-orbitally injected (0.2 ml/mouse) to 8 weeks old recipient mice. Blood samples were collected retro-orbitally with a heparin-containing tube at various time points after transfusion. 500 μl saline was added to the blood samples. Platelets were isolated by centrifugation at 1,000 rpm for 3 min and analyzed by flow cytometry. Ratio of GFP positive platelets in total platelets in fresh PRP group at 5min after transfusion was set to 100% (survival 100%). Survival of the transfused platelets was normalized to the survival of transfused fresh platelets at 5 min after transfusion.

### Fibrinogen binding to platelets

Washed platelets (3 × 10^8^/ml) after incubated at RT or 4°C for 24 h, were added with 1 mM CaCl_2_ and 2 μl FITC-labeled fibrinogen and incubated at 22°C for 30 min with gentle shaking. Fibrinogen binding to cells was analyzed by flow cytometry.

### Measurement of serotonin in platelet supernatant

Washed platelets (5 × 10^8^/ml 500 μl) from C57BL/6J mice or healthy donors were pre-incubated with vehicle, EGTA, or RGDS at RT for 15 min and then stored at RT or 4°C for 24h. The supernatant was collected by centrifuging at 12,000 rpm for 1 min and 0.1N HCl was added. 1M TCA was added to the supernatant. The TCA extract was transferred to a tube containing 2 ml of 0.05% O-phalaldehyde after centrifuging at 12,000 rpm for 2 min at room temperature. Samples were boiled for 10 min, cooled on ice, and then washed twice with chloroform. Serotonin release was measured using an MDS fluorescence spectrophotometer (MDS Analytical Technologies, Sunnyvale, CA) with an excitation wavelength of 360 nm and an emission wavelength of 475 nm.

### Measurement of PF4 in platelet supernatant

Washed platelets (3 × 10^8^/ml) from C57BL/6J mice were pre-incubated with vehicle, EGTA, or RGDS at RT for 15 min and then stored at RT or 4°C for 24 h. After incubation, supernatant of washed platelets was collected by centrifuging at 3,000 rpm for 4 min. The amount of PF4 in platelet supernatant was measured using the anti-mouse ELISA kit from R & D Systems (Minneapolis, MN) following the manufacturer’s instruction.

### Measurement of VWF secretion by Western blot

Washed platelets (3 × 10^8^/ml) from C57BL/6J mice were stored at 4°C for 24 h. After incubation, supernatant of washed platelets was collected by centrifuging at 3,000 rpm for 4 min. VWF in supernatant was detected by Western blot with a rabbit polyclonal antibody against VWF ((Dako Agilent Technologies, Santa Clara, CA).

### Western blot analysis of Src phosphorylation

Washed platelets (3 × 10^8^/ml) were stored at 4°C or RT for 0.5 h, 2 h, 4 h, and 24 h in pH SAFE Mini bags and solubilized with 2 x SDS-PAGE sample buffer. Platelet lysates were analyzed by SDS-PAGE on 4∼15% gradient gels and immunoblotted using a polyclonal anti-phosphorylated Src residue Tyr416.

### Platelet spreading on fibrinogen

Washed platelets (2.5 × 10^7^ /ml) from C57BL/6J mice were resuspended in Tyrode’s solution and were pretreated with 100 μm EGTA or vehicle at RT for 10 min, followed by the addition of CaCl_2_ or control buffer and incubated at RT for 10 min. Chamber slides (Lab-Tek, Nunc, Thermo-Fisher Scientific, Rochester, NY) were coated with 50 μg/ml fibrinogen and blocked with 5% bovine serum albumin (BSA) in phosphate-buffered saline (PBS). Platelets were added to wells (100 μl/well) and allowed to spread for 2h at 37 °C. Slides were rinsed to detach non-adherent platelets and fixed with 1% paraformaldehyde. Samples were blocked with 5% BSA in PBS and incubated with Rhodamine-conjugated phalloidin. Images were collected with a Leica DM IRB microscope using 60 x objective (Leica Microsystems, Wetzlar, Germany), and platelet size was quantified using Image J. Quantification of area was normalized to platelets without treatment. Statistical analysis performed using unpaired Student t test.

### Measurement of mouse tail-bleeding times and blood loss

Mouse PRP from C57BL/6J mice was stored at RT or 4°C for 48 hours in the presence or absence of 2 mM EGTA. PRP in the presence of EGTA was supplemented with 2 mM CaCl_2_ before transfusion. 1 × 10^9^ platelets in 200 μl PRP were injected retro-orbitally into 7∼8 week-old mice. After 2 hours, the mice were anesthetized by inhalation of 2∼5% isoflurane in 100% oxygen using anesthesia equipment. The distal portion of the tail (1 mm) was amputated with a scalpel, and the tail was immersed in 0.15 M NaCl at 37°C as previously described. Time to stable cessation of the bleeding was defined as the time where no rebleeding for longer than 2 min was recorded. Statistical analysis was performed using nonparametric Mann-Whitney test. To measure blood loss volume, any blood collected from tail transection was frozen at −80°C overnight. After thawing the following day, 10 ml of deionized water was added to further induce hemolysis. Aliquots of each sample were analyzed via spectrophotometry (SpectraMax Plus384; Molecular Devices, Sunnyvale, CA) and diluted further (1:5, 1:10, or 1:20) if necessary. The resulting OD490 nm values (% T) were compared against a standard curve to estimate the blood volume lost.

### Western blot analysis of caspase-3 cleavage

Washed mouse platelets were stored at 4°C for 4 h. Platelet lysates were analyzed by SDS-PAGE on 4∼15% gradient gels and immunoblotted using a rabbit polyclonal antibody that recognizes both caspase-3 and an active fragment of caspase-3.

## Acknowledgments

B.X. is supported by AHA Great Rivers Affiliate Scientist Development Grant. C.W. is supported by an K99/R00 grant (K99HL145117; NHLBI). Z.L. is supported by NIH/NHLBI R01 HL123927, R01 HL142640, and R01HL146744, and NIH/NIGMS R01 GM132443. We thank Dr. Richard H. Aster for critical reading of the manuscript and suggestions.

## Contributions

B.X., G.Z., and Z.L. designed and performed most of the experiments, analyzed and interpreted the data, and wrote the manuscript. Y.Z. and C.W. performed and helped with the *in vivo* experiments. S.J. and S.W.W. performed and helped to analyze platelet secretion. A.J.M, J.W., S.S.S., and S.W.W. provided conceptual advice and helped to design experiments. All authors critically analyzed data, and edited and approved the manuscript.

