## Supplemental figures for "Calcium ion chelation preserves platelet function during cold storage"

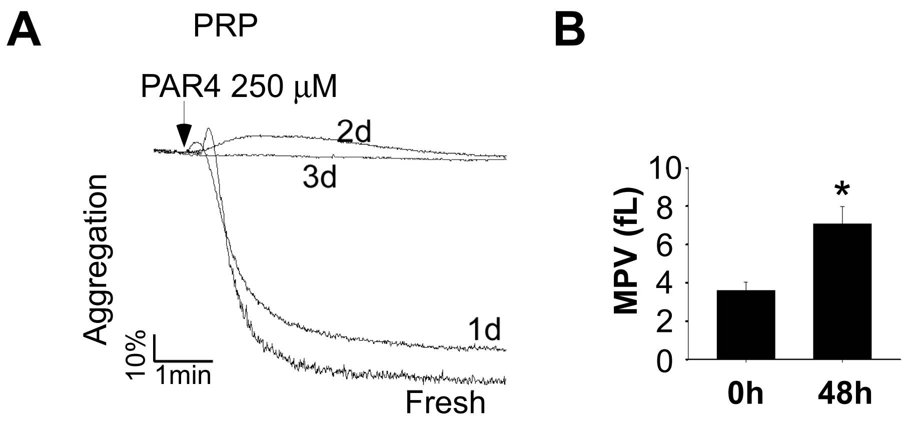


**Supplementary Figure 1. Room temperature storage caused lesions to platelets.** (a) PRP from C57BL/6J mice was stored at RT for various lengths of time. Platelet aggregation was induced by addition of the thrombin receptor agonist PAR4. (b) Platelet size after RT storage was measured by a HEMAVET HV950FS multispecies hematology analyzer.

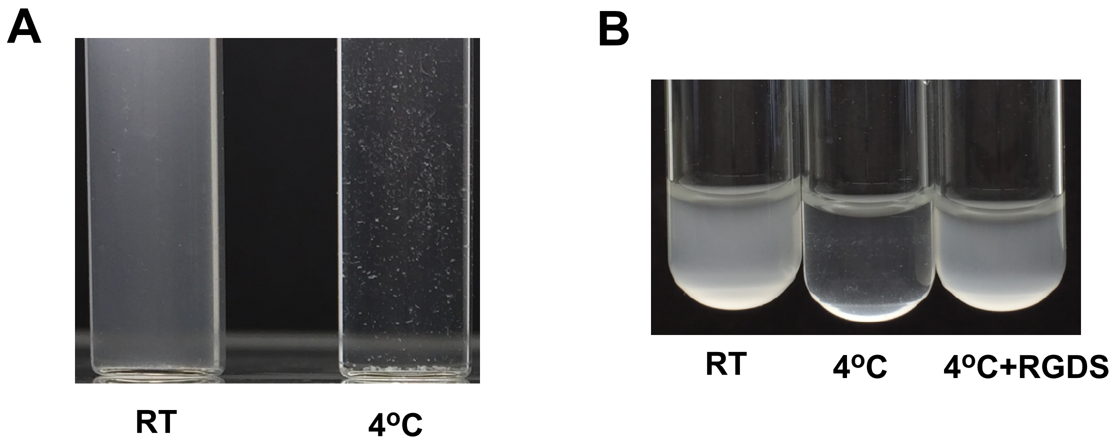


**Supplementary Figure 2. Cold induced platelet aggregation.** (a) Washed platelets from C57BL/6J mice (3 x $10^8$/ml) were stored at RT or 4°C for 24h and transferred to a new glass tube for taking image. A representative image is shown. (b) Washed platelets (3 x $10^8$/ml) from C57BL/6J mice were preincubated with vehicle or RGDS (1mM) for 15 min at RT and then stored at RT or 4°C for 24 h. After incubation, platelets were transferred to a new glass tube for taking image.

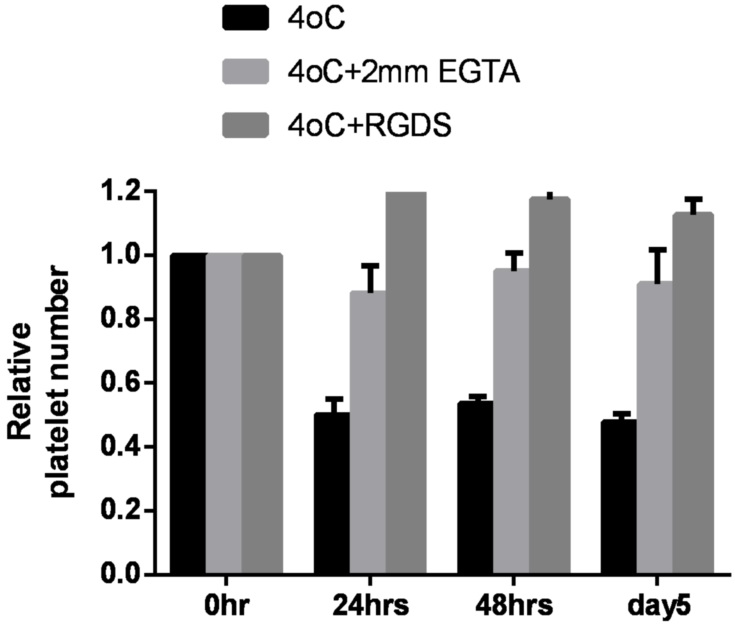


**Supplementary Figure 3. Platelet counts decreased after cold storage.** Washed platelets from healthy donors were stored at RT or 4°C in the presence of absence of EGTA (2 mM) or RGDS (1 mM). Platelet counts were monitored at various time points.

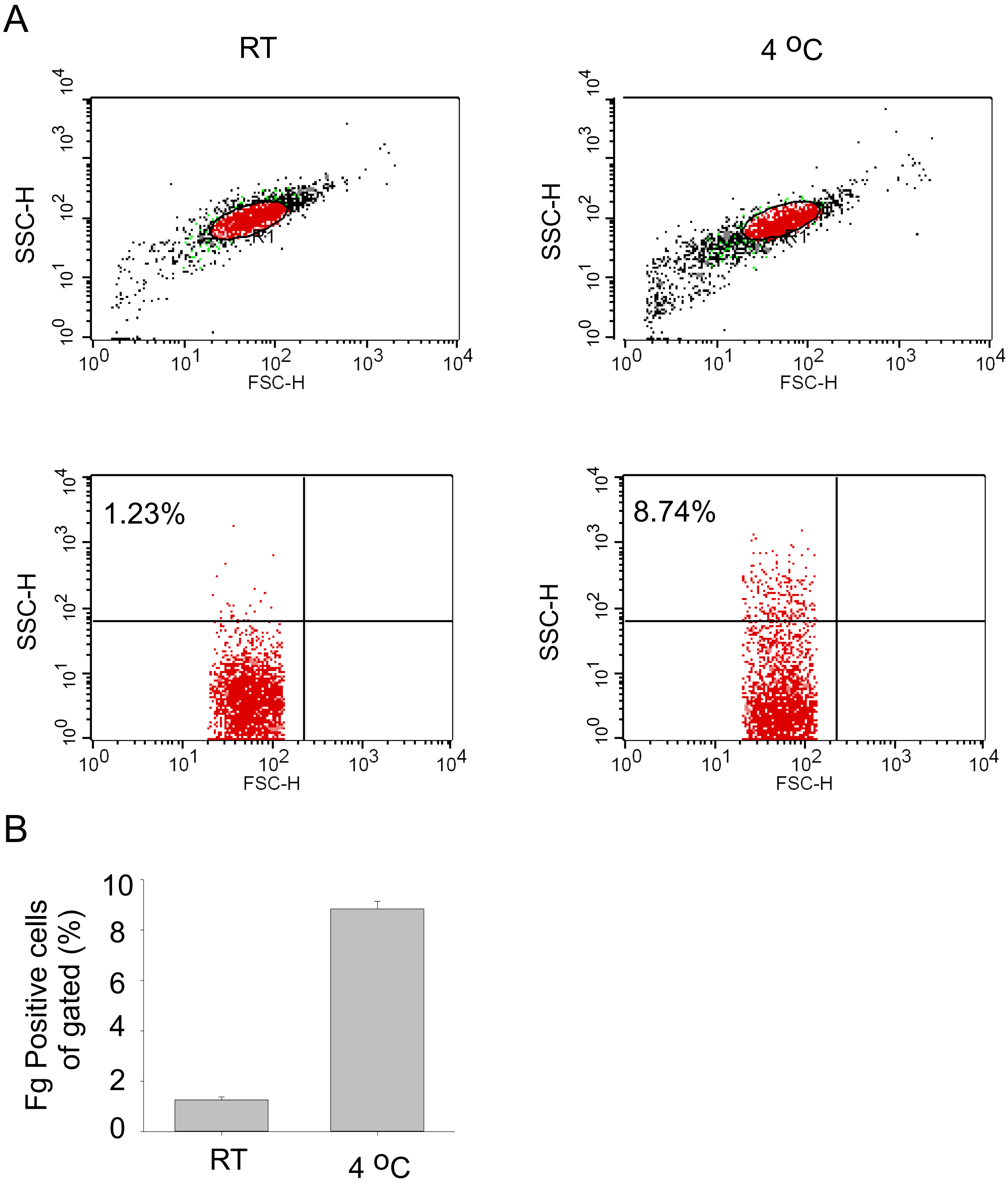


**Supplementary Figure 4. Fibrinogen binding to platelets was increased after cold storage.** (a) Washed human platelets were incubated at RT or 4 °C for 24 hours. FITC-labled fibrinogen was added to platelets and incubated at RT for 30 min and then analyzed by flow cytometry. (b) Fibrinogen positive cells of the gated cells (n-4, $p<0.01$ by unpaired *t-test*).

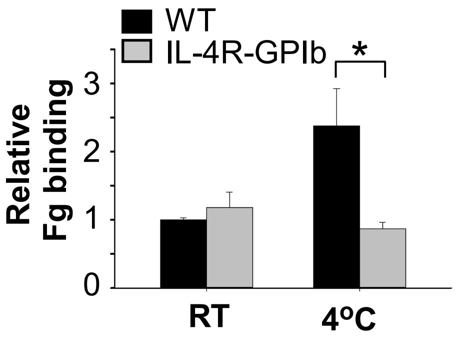


**Supplementary Figure 5. GPIb was involved in cold-induced activation of platelet integrin αIIbβ3.** Washed platelets from wild-type mice or IL-4R-GPIb mice were stored at RT or 4oC for 24 h. Integrin αIIbβ3 activation was analyzed by FITC-labeled fibrinogen binding using flow cytometry. Quantification of fibrinogen (Fg) binding relative to fibrinogen binding of wild-type platelets at RT for 24 h was shown. The statistical differences were examined by Student t test. * $p<0.05$.

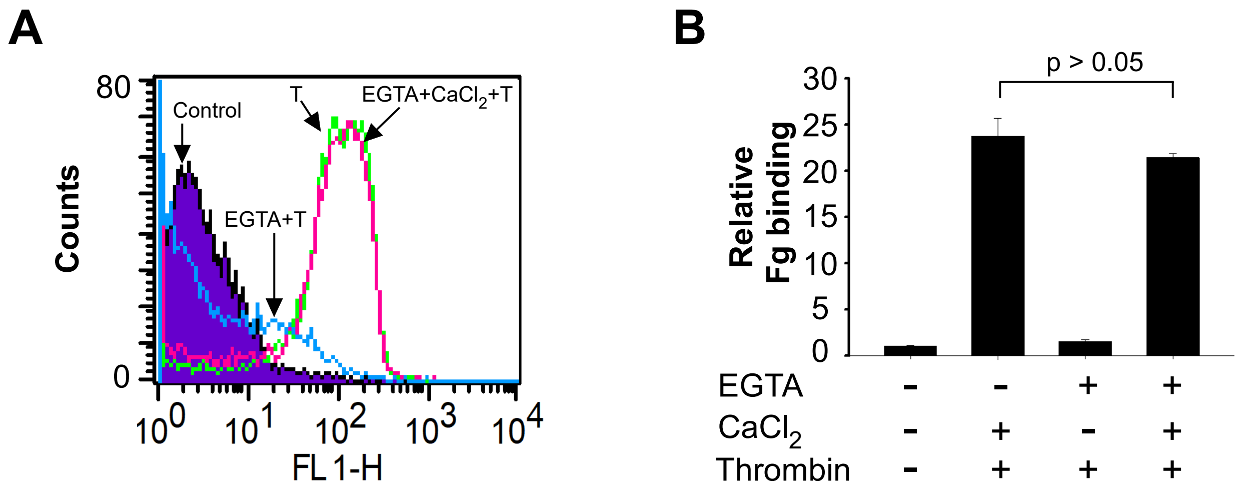


**Supplementary Figure 6. Inhibition of platelet integrin αIIbβ3 activation by EGTA was reversible.** (**a-b**) Washed platelets (3 x $10^8$/ml) were preincubated with 100uM EGTA or vehicle at RT for 10 min and then added with or without $CaCl_2$ at RT for 3 min. After treatment, platelets were stimulated with 0.05U/ml thrombin (T) in the presence of FITC-labeled fibrinogen at RT for 30 min. Fibrinogen binding was analyzed using flow cytometry. A representative flow cytometry plots (**a**) and quantification of fibrinogen (Fg) binding relative to vehicle-treated control (**b**) were shown.

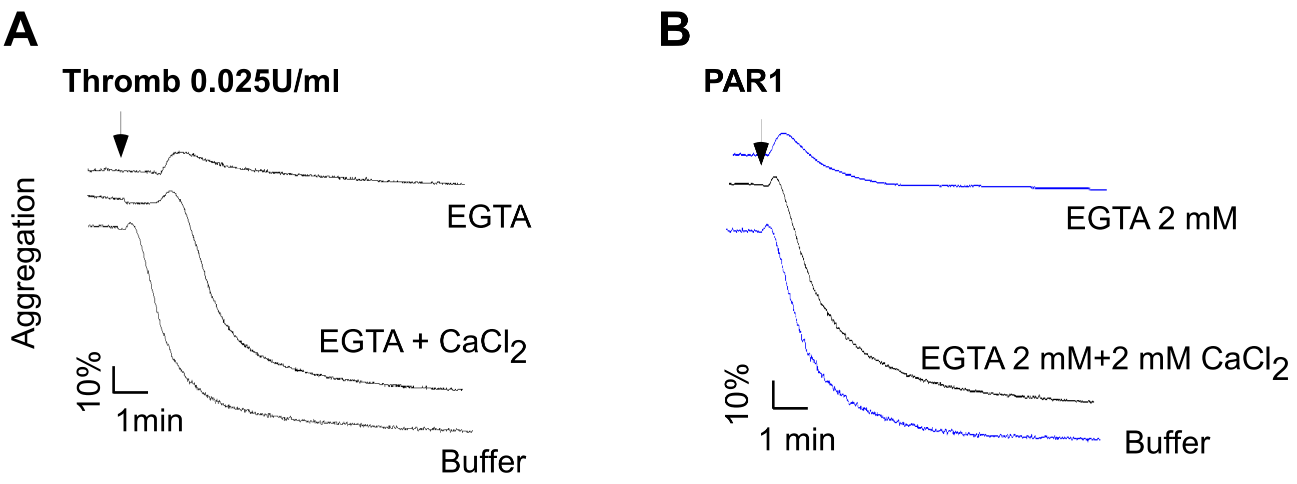


**Supplementary Figure 7. EGTA reversibly inhibited platelet aggregation.** (a) Washed platelets from C57BL/6J mice (3 x $10^8$/ml) were treated with or without 100 µM EGTA for 10 min at room temperature. Then platelets were added with or without 1mM $CaCl_2$ and stimulated with 0.025U/ml Thrombin in a lumi-aggregometer at 37°C. (b) PRP from a healthy donor was incubated with 2 mM EGTA for 5 min, and then added with 2 mM $CaCl_2$ or buffer. Platelet aggregation was induced by addition of 2.5 µM PAR1 peptide.

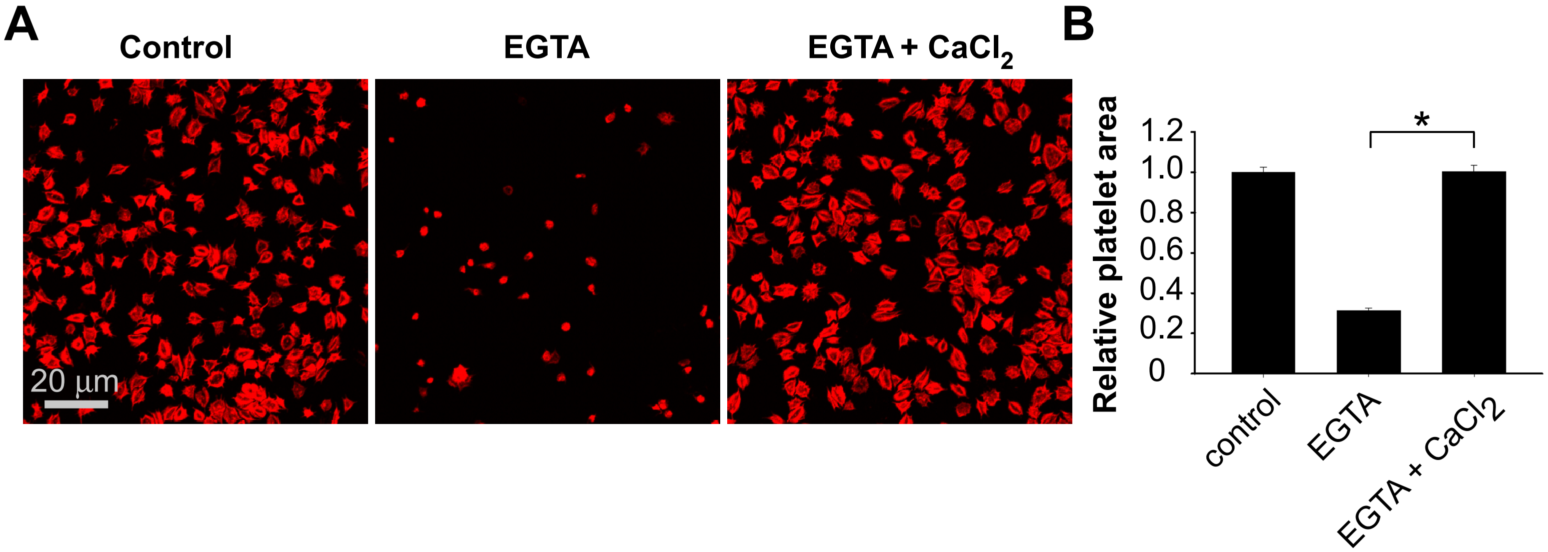


**Supplementary Figure 8. Inhibition of platelet spreading by EGTA was reversible.** (**a-b**) Washed platelets from C57BL/6 mice (2.5 x $10^7$/ml) were pretreated with 100uM EGTA or vehicle at RT and then added with or without $CaCl_2$ for 10min. Platelets were added to 50μg/ml fibrinogen-coated slides for 2h at 37°C and were fixed with paraformaldehyde. Platelets were labeled with Rhodamine-Phalloidin and photographed using a fluorescence microscope (**a**). Platelet area was quantified and normalized to platelets without treatment (**b**) (mean±SEM; n=117–246; *P<0.0001).

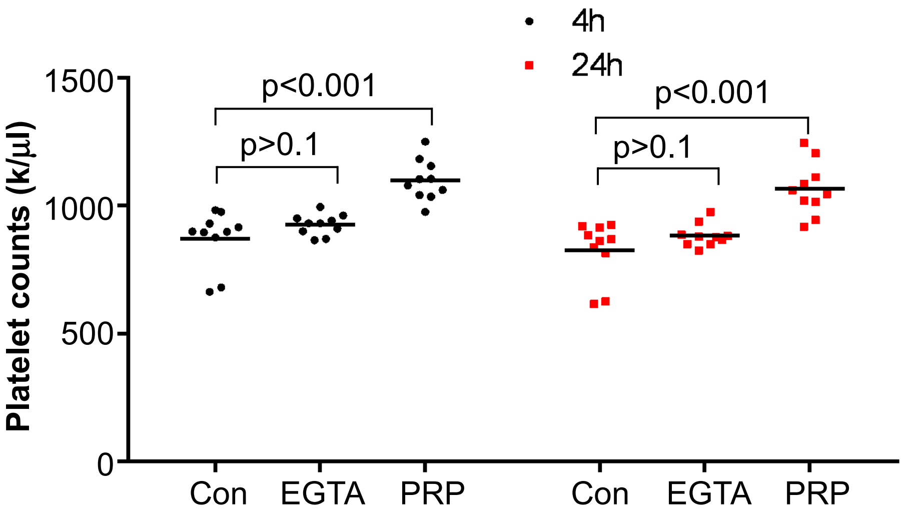


**Supplementary Figure 9. Transfusion of EGTA-treated platelets or EGTA alone did not cause thrombocytopenia**. (a) PRP (1.25 x 109/ml) from C57BL/6J mice were added with EGTA (2 mM) and incubated at RT for 15 min. C57BL/6J mice were injected with 0.2 ml of PRP+EGTA or 0.2 ml of 2 mM EGTA in saline per mouse. After 14 days, the mice were injected with the same amount of PRP+EGTA or EGTA in saline. Platelet counts were monitored with a HEMAVET HV950FS multispecies hematology analyzer at 4 hours and 24 hours after the second transfusion of PRP/EGTA or EGTA/saline. The statistical differences were examined by unpaired Student t test.
